# Hypoxia responses in arginase 2 deficient mice enhance cardiovascular health

**DOI:** 10.1101/2025.02.20.639297

**Authors:** Weiling Xu, Kewal Asosingh, Allison J. Janocha, Evan Madden, Nicholas Wanner, Dylan Trotter, Michael V. Novotny, Anny Mulya, Samar Farha, Serpil C. Erzurum

**Author notes:** Address correspondence to Serpil C. Erzurum, M.D., Cleveland Clinic, 9500 Euclid Avenue, NC22 Cleveland, Ohio 44195, USA.

## Abstract

**RATIONALE:** Physiological responses to hypoxia involve adaptations in the hematopoietic and cardiovascular systems, which work together to ensure adequate oxygen delivery to tissues for energy production. The arginine/nitric oxide (NO) pathway regulates both systems through its effects on erythropoiesis and vasodilation. In Tibetan populations native to high-altitude hypoxia, increased NO production from arginine and decreased arginine metabolism by arginase contribute to these adaptive mechanisms. These metabolic changes enhance tissue oxygen delivery and reduce the risk of hypoxic pulmonary hypertension. Here, we hypothesize that genetic deletion of mitochondrial arginase 2 (*Arg2*) in mice will enhance cardiovascular effects and mitigate hypoxia-induced pulmonary hypertension.

**METHODS:** Complete blood counts, bone marrow erythroid differentiation, plasma arginine and NO (measured as nitrite), right ventricular systolic pressure (RVSP), heart rate, heart weight, and blood pressure were measured in wild-type (WT) and *Arg2* knockout (*Arg2*KO) mice exposed to short-term (6, 12, 48, or 72 hours) or long-term (3 weeks) hypoxia.

**RESULTS:** Under normoxic conditions, *Arg2*KO and WT mice exhibit similar RBC counts, hemoglobin levels, hematocrit, heart rate, systolic and diastolic blood pressures, and heart weight (all *P* > 0.05). WT mice increase erythropoiesis at 12 hours of hypoxia, including proerythroblasts (stage I, *P* = 0.004), polychromatic erythroblasts (stage III, *P* = 0.0004), and orthochromatic erythroblasts (stage IV, *P* = 0.03), but *Arg2*KO mice do not increase erythropoiesis. After 48 hours of hypoxia, *Arg2*KO mice increase proerythroblasts (stage I, *P* = 0.0008), but levels remain significantly lower than in WT mice. Plasma arginine and NO levels increase under hypoxia. NO levels peak at 12 hours of hypoxia in WT mice, then decline rapidly. In contrast, NO levels in *Arg2*KO mice are higher than in WT mice, with sustained elevations at 48 hours of hypoxia (*P* = 0.03). *Arg2*KO mice have significantly higher plasma arginine levels than WT at 6, 12, and 72 hours of hypoxia (all *P* < 0.05). Under chronic hypoxia, *Arg2*KO and WT mice show similar RBC counts, hemoglobin levels, hematocrit, and NO levels. Unlike WT, *Arg2*KO mice do not increase RVSP (*P* = 0.4) and have lower mean arterial (*P* = 0.03) and diastolic blood pressures (*P* = 0.01), as well as much lower heart rates (*P* < 0.0001). Additionally, small blood vessels increase in lungs of *Arg2*KO mice (CD31, *P* = 0.02; vWF, *P* = 0.6).

**CONCLUSIONS:** Arginine metabolism in the mitochondria plays a key role in modulating adaptive responses to hypoxia. Deletion of *Arg2* results in delayed erythropoiesis under acute hypoxia, but better cardiovascular health, as indicated by higher levels of nitrite and arginine, and lower RVSP, blood pressure, and heart rate with chronic hypoxia.

## Introduction

Major physiological responses to enhance oxygen delivery include vascular adaptations (including increased nitric oxide (NO) production, blood vessel growth, and vasodilation) to increase blood flow; hematopoietic adaptations (erythropoiesis) to improve oxygen-carrying capacity; and cardiac adaptations (such as alterations in heart rate, blood pressure, and cardiac output) (1–15). High-altitude populations utilize these biological adaptations to optimize oxygen delivery: vascular adaptations to increase blood flow in Tibetans and Ethiopians, and hematological adaptations to enhance oxygen-carrying capacity in Andeans (1–15). Tibetan populations native to high-altitude hypoxic environments exhibit low arginase activity and high arginine bioavailability, which promote greater NO production and cGMP activation, resulting in vasodilation and increased blood flow (1, 2, 6, 15). These metabolic adaptations enhance tissue oxygen delivery and reduce the risk of hypoxic pulmonary hypertension in such environments.

The arginine/nitric oxide (NO) pathway regulates both hematopoietic and cardiovascular systems through effects on erythropoiesis and vasodilation (16–22). Arginine, a semi-essential amino acid, is the substrate for NO synthase and arginases (ARG). Endothelial nitric oxide synthase (eNOS), the predominant NOS isoform in the pulmonary vasculature, converts arginine to NO and citrulline. NO is a potent vasodilator that is deficient in pulmonary hypertension (17–20, 23). Under certain *in vivo* conditions, arginine bioavailability may limit the production of NO, e.g., arginine utilized by other enzymes, such as ARG (24–26). ARG catabolizes arginine to ornithine and urea (26). ARG1 is present exclusively in the cytosol of hepatic cells as part of the urea cycle, but ARG2, encoded by nuclear DNA, is found in mitochondria of many tissues without a functioning complete urea cycle, including the lungs and heart. Loss of NO production in pulmonary arterial hypertension is associated with greater mitochondrial arginine metabolism *via* ARG2 (21, 26). Here, we hypothesize that the genetic deletion of mitochondrial arginase 2 (*Arg2*) in mice will be sufficient to improve oxygen delivery through hematologic and cardiovascular effects, and to mitigate the development of pulmonary hypertension. The study of acute and chronic hypoxia responses provides a comprehensive framework to study the progression of hypoxia-induced pulmonary hypertension from acute hematopoietic responses to chronic cardiopulmonary responses.

## Methods

### Wild-type (WT) and arginase 2 knockout (Arg2KO) mice

Male mice on a C57BL/6J background at the age of 11-12 weeks were used in this study. WT mice were purchased from the Jackson Laboratory (Bar Harbor, ME) and *Arg*2KO were donated by W.E. O’Brien (Baylor College of Medicine, Houston, TX) (27).

*Hypoxia model.* WT and *Arg2*KO mice were exposed to short-term (6-, 12-, 48-, or 72-hours) or long-term hypoxia (3-weeks) or kept under normoxia as described (28, 29). A chamber equipped with passive ventilation openings (Small Chamber 38 cm × 51 cm × 51 cm; BioSpherix, Redfield, NY) and connected to a secured liquid nitrogen tank to control the oxygen levels within the chamber was utilized for hypoxia experiments. The chamber’s atmospheric conditions were established at standard atmospheric pressure, featuring 10% oxygen, which was regulated by an oxygen sensor (ProOx model 360, BioSpherix). The calibration of the oxygen sensor was performed in accordance with the manufacturer’s guidelines. A single set point of 10% oxygen was established in the hypoxia chamber, with an alarm triggered at 11% to monitor the oxygen levels.

### NO (measured as NOx)

NOx [nitrite (NO_2_ ^-^) and nitrate (NO_3_ ^-^)] concentrations were determined by using the Eicom NOx Analyzer ENO-30 HPLC system (Amuza Inc. San Diego, CA). Murine serum samples were deproteinated with an equal volume of methanol, chilled, then pelleted to remove protein by centrifuging at 18,000 x*g* at 4°C for 30 mins (Sorvall Legend Micro 21R Centrifuge, Thermo Fisher Scientific, Waltham, MA). Supernatants were loaded into a 96-well plate. 10 µL was injected into the ENO-30 using the Eicom AS-700 Insight Autosampler. Respective peaks for NO_2_ ^-^ and NO_3_ ^-^ were monitored using Eicom Envision SPC-700 software. The concentrations of NO_2_ ^-^ and NO_3_ ^-^ were calculated by comparing the peak areas of samples with peak areas of known NO_2_ ^-^ and NO_3_ ^-^ standards. Samples were run in duplicate.

### Amino acid analysis

Plasma arginine was measured by LC/MS/MS methodology (30). Briefly, 20 μL plasma was methanol-extracted in the presence of 20 μM of phenylalanine-d5 (internal standard). 5 μL of methanol extracted samples were injected into Vanquish UHPLC – TSA Quantiva (Thermo Fisher Scientific) and separated with C18 LC column (Gemini 3 μm, 2×150 mm) (Phenomenex, Torrance, CA) with flow rate of 0.3 mL/min. Mobile phase for amino acid separation were [A] 0.2%v/v acetic acid and 10 mM ammonium acetate and [B] methanol/acetonitrile (1:1, v/v), 0.2%v/v acetic acid and 10 mM ammonium acetate. Selected reaction monitoring (SRM) transition was detected at electrospray ionization (ESI) positive ionization [SRM transition, ESI positive ionization, arginine 175 > 116]. Data acquisition and interpretation was performed with Xcalibur v4.1 software (Thermo Fisher Scientific).

### Flow cytometry for bone marrow erythropoiesis

Bone marrow was flushed from the long bones of the hind legs with phosphate buffered saline (PBS) before filtering through a 30 µm filter. Fc receptors were blocked in 50 μL of CD16/32 antibody (eBioscience 14-0161-85, clone: 93, 0.5 µg/50µL) diluted in 1% bovine serum albumin (BSA) for 15 minutes, followed by co-incubation with the following cell surface antibodies in a final volume of 100 µL for 30 minutes at 4°C: D45-APC (eBioscience 17-0451-82, clone: 30-F11; 0.2 µg/100µL), CD11b-APC (eBioscience 17-0112-81, M1/70; 3.1 ng/100µL), Ly-6G-APC (eBioscience 17-9668-80, clone: 1A8-Ly6g; 0.13 µg/100µL), TER119-PE (eBioscience 12-5921-81, clone: TER-119; 0.13 µg/100µL), CD44-FITC (eBioscience 11-0441-81, clone: IM7; 0.25 µg/100µL) (1,2). After incubation, the cells were washed with 1% BSA in PBS and resuspended in 500 µL of 1% BSA with 7-AAD (BD 559925; 1.25 µg/500µL). Data were acquired using the Sony ID7000 spectral flow cytometer equipped with 6 lasers (320nm, 355nm, 405nm, 488nm, 561nm, 637nm) collecting 3 million events per sample. Daily quality control of the cytometer was performed to ensure instrument consistency and track performance. Spectral unmixing was done using single color controls in the Sony ID7000 analysis software before loading unmixed FCS files into FlowJo (10.9.0) for gating. Fluorescence minus one (FMO) sample were used as gating controls. Aggregates, cell debris, and dead cells were removed from the analysis, then APC positive cells (CD45+, CD11b+, Ly-6G+) were excluded. All TER119 positive cells were gated, then proerythroblasts were defined as CD44hi+/TER119lo+. Basophilic erythroblasts, polychromatic erythroblasts, and orthochromatic erythroblasts were defined as TER119hi+ and selected based on decreasing size and CD44 expression. Within each experiment, gates were set on concatenated .fcs files for samples under normoxia, starting with stage I (proerythroblasts) to obtain the expected ratio of 1:2:4:8 (stages I:II:III:IV). Individual samples were deconvoluted after gating for the expected ratio (31, 32).

### Complete blood count (CBC)

CBC was measured as using a fully automated Advia 120 Hematology system (Siemens, Cary, NC)

### Right ventricular systolic pressure (RVSP)

RVSP, heart rate, heart weight, and blood pressure were collected as described previously (28, 29).

### Lung tissue harvest and immunohistochemistry

The lung vascular bed was perfused with warm PBS via the right heart to flush out remaining blood cells. The dissected left lobe was fixed in 10% formalin and processed for paraffin embedding and tissue sectioning. Left lung tissue sections were stained with CD31 and von Willebrand Factor (vWF) staining, and quantification was performed as described previously (28). Rabbit anti-CD31 (Santa Cruz Biotechnology, Dallas, TX) at a 1:50 dilution and anti-vWF (Dako, Carpinteria, CA) at a 1:200 dilution were used, and the slides were counterstained with hematoxylin and bluing.

*Statistics.* Analyses were conducted with JMP Pro 18 (SAS Institute, Cary, NC). Data were summarized as mean ± SD. For comparisons of two groups, Student’s t-test or Wilcoxon test was used as appropriate. For three or more groups, analysis of variance (ANOVA) was used. The level of significance for *P* was chosen at 0.05.

### Study approval

All animal experiments were approved by the Cleveland Clinic IACUC (Cleveland, Ohio).

## Results

### Under normoxia, lower erythroid progenitors and higher plasma arginine in lung of *Arg*2KO mice as compared with WT mice

Under normoxia, *Arg*2KO mice trend to higher plasma NO (*P* = 0.06) (Table 1) (Fig. 1A), and have higher plasma arginine (*P* < 0.0001) compared to WT mice (Table 1) (Fig. 1B) similar to a previous report (27). *Arg*2KO and WT mice have similar heart rate, systolic and diastolic blood pressures, and heart weight (all *P* > 0.05) (Table 2), suggesting that hematopoietic and cardiovascular systems are not affected by the changes in arginine metabolism in *Arg2*KO mice under normoxia.

**Fig. 1.**
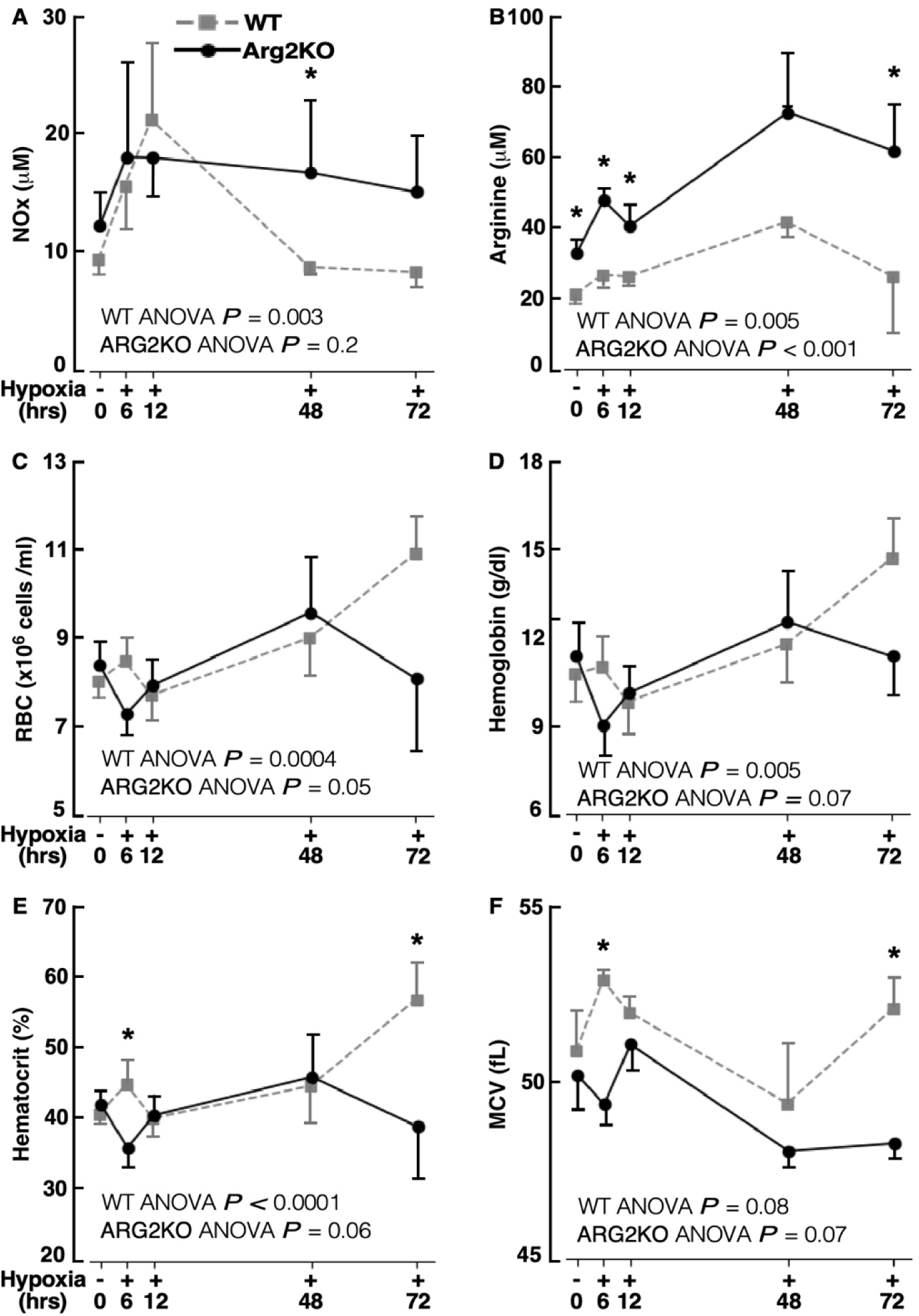
Amount of NO products (NOx), arginine, red blood cells (RBC), hemoglobin, hematocrit, and mean corpuscular volume (MCV) in *Arg2*KO and WT mice under short-term (6-, 12-, 48-, or 72-hours) hypoxia or kept under normoxia. [A – B] Plasma NOx (μM) (**A**) and arginine amount (μM) (**B**) in *Arg*2KO *vs.* WT mice. **[C – F]** Blood RBC (**C**), hemoglobin (**D**), hematocrit (**E**), and MCV (**F**) in complete blood count (CBC) of *Arg*2KO and WT mice. *n* ≥ 3 for each group. \**P* < 0.05 for *Arg2KO vs*. WT at the corresponding time point.

**Table 1.** Plasma arginine and NO levels, erythropoietic response, and complete blood counts of *Arg*2KO and WT mice under short-term hypoxia or kept under normoxia.

| Characteristics | WT (*n* ≥ 3) |  |  |  |  |  | Arg2KO (*n* ≥ 3) |  |  |  |  |  | *P* value*** |
| --- | --- | --- | --- | --- | --- | --- | --- | --- | --- | --- | --- | --- | --- |
|  | Normoxia | Hypoxia |  |  |  | *P* value* | Normoxia | Hypoxia |  |  |  | *P* value** |  |
|  |  | 6h | 12h | 48h | 72h |  |  | 6h | 12h | 48h | 72h |  |  |
| Plasma |  |  |  |  |  |  |  |  |  |  |  |  |  |
| NOx (μM) | 9.5(2.2) | 15.8(3.5) | 21.3(7.9) | **8.8(0.9)** | 8.4(2.0) | **0.003** | 12.3(4.6) | 18.2(6.9) | 18.1(4.6) | **16.8(5.9)** | 15.2(7.0) | 0.2 | 0.06 |
| Arginine (μM) | **21.0(3.6)** | **26.7(5.5)** | **26.2(3.3)** | 41.5(4.8) | **26.3(14.60)** | **0.0005** | **32.8(5.6)** | **47.7(3.5)** | **40.4(7.4)** | 72.5(22.8) | **61.7(11.7)** | **<0.001** | **<.0001** |
| Blood |  |  |  |  |  |  |  |  |  |  |  |  |  |
| RBC (x10<sup>6</sup> cells/mL) | 8.1(1.0) | 8.5(0.6) | 7.7(1.0) | 9.0(1.1) | 10.9(0.9) | **0.0004** | 8.4(1.0) | 7.3(0.6) | 8.0(1.0) | 9.6(1.3) | 8.1(1.6) | 0.05 | 0.3 |
| Hemoglobin (g/dl) | 10.7(2.0) | 11.0(1.0) | 9.8(1.8) | 11.8(1.3) | 14.7(1.5) | **0.005** | 11.4(2.1) | 9.0(1.0) | 10.1(1.5) | 12.5(1.7) | 11.3(1.5) | 0.07 | 0.4 |
| Hematocrit (%) | 40.9(3.6) | **45.0(3.6)** | 40.2(4.9) | 44.8(6.7) | **57.0(6.2)** | **<.0001** | 42.1(3.7) | **36.0(3.0)** | 40.6(4.2) | 46.0(6.2) | **39.0(7.5)** | 0.06 | 0.3 |
| MCV (fL) | 50.9(2.5) | **52.9(0.4)** | 52.0(0.6) | 49.4(1.8) | **52.1(1.4)** | 0.08 | 50.2(1.8) | **49.4(0.5)** | 51.1(1.3) | 48.1(0.5) | **48.3(0.4)** | 0.07 | 0.4 |
| MCH (pg) | 13.3(0.9) | 13.0(0.5) | 12.7(0.6) | 13.2(0.4) | 13.4(0.1) | 0.3 | 13.5(1.0) | 12.0(0.5) | 12.8(0.3) | 13.1(0.3) | 13.8(1.5) | **0.04** | 0.7 |
| RDW (%) | 12.5(1.0) | 12.4(0.4) | 12.5(0.8) | 14.7(0.9) | 14.2(0.7) | **0.0002** | 12.2(0.3) | 12.6(0.2) | 12.3(0.6) | 14.2(0.4) | 14.9(0.6) | **<.0001** | 0.4 |
| Platelets (x10<sup>6</sup> cells/mL) | 0.70(0.25) | 0.69(0.16) | 0.78(0.22) | 0.93(0.28) | 1.04(0.28) | 0.16 | 0.83(0.11) | 0.75(0.10) | 0.75(0.26) | 0.63(0.32) | 1.14(0.53) | 0.09 | 0.09 |
| WBC (x10<sup>3</sup> cells/mL) | 2.4(1.2) | **2.7(0.6)** | 1.6(0.7) | **5.8(1.9)** | **4.6(0.9)** | **<.0001** | 2.5(1.1) | **1.3(0.1)** | 2.4(1.2) | **2.5(0.8)** | **2.3(0.6)** | 0.4 | 0.7 |
| Bone Marrow<sup>#</sup> |  |  |  |  |  |  |  |  |  |  |  |  |  |
| Stage I | 1.3(0.4) | 1.0(0.2) | **2.4(0.7)** | **1.1(0.1)** | 3.1(0.2) | **<.0001** | 0.9(0.1) | 0.9(0.2) | **0.8(0.2)** | **1.8(0.2)** | 2.7(0.4) | **<.0001** | 0.12 |
| Stage II | **2.4(0.4)** | 1.9(0.2) | 1.9(0.4) | 2.6(0.3) | 3.6(0.3) | **<.0001** | **1.6(0.2)** | 2.3(0.3) | 1.5(0.1) | 2.9(0.3) | 3.8(0.7) | **<.0001** | **0.01** |
| Stage III | **5.3(0.8)** | 3.7(0.6) | **4.8(0.5)** | 5.0(0.9) | 6.4(0.4) | **0.003** | **3.5(0.3)** | 4.7(1.1) | **3.0(0.2)** | 5.6(0.4) | 6.8(0.5) | **<.0001** | **0.005** |
| Stage IV | **10.4(1.5)** | 7.8(1.2) | **10.1(1.5)** | 11.5(1.8) | 11.4(0.6) | **0.03** | **7.9(0.6)** | 9.1(2.7) | **8.7(0.7)** | 13.1(1.1) | 14.0(1.5) | **<.0001** | **0.01** |

Mean (SD).
Bold values indicate *P* < 0.05 for *Arg2*KO vs. WT at the corresponding time point.
\**P* value for ANOVA of normoxia and hypoxia in WT; \*\**P* value for ANOVA of normoxia and hypoxia in *Arg2*KO; \*\*\**P* value for *Arg2*KO vs. WT under normoxia.
<sup>#</sup>Proportion in erythropoietic stage of proerythroblasts (stage I), basophilic erythroblasts (stage II), polychromatic erythroblasts (stage III), and orthochromatic erythroblasts (stage IV) was determined based on expression of TER119, CD44, and size.
Definition of abbreviations: *Arg2*KO, genetic deletion of *Arg2* or *Arg2*<sup>-/-</sup> mice; MCH, mean corpuscular hemoglobin; MCV, mean corpuscular volume; NOx, NO products; RBC, red blood cells; RDW, red blood cell distribution width; WBC, white blood cells; WT, wildtype.

**Table 2.**
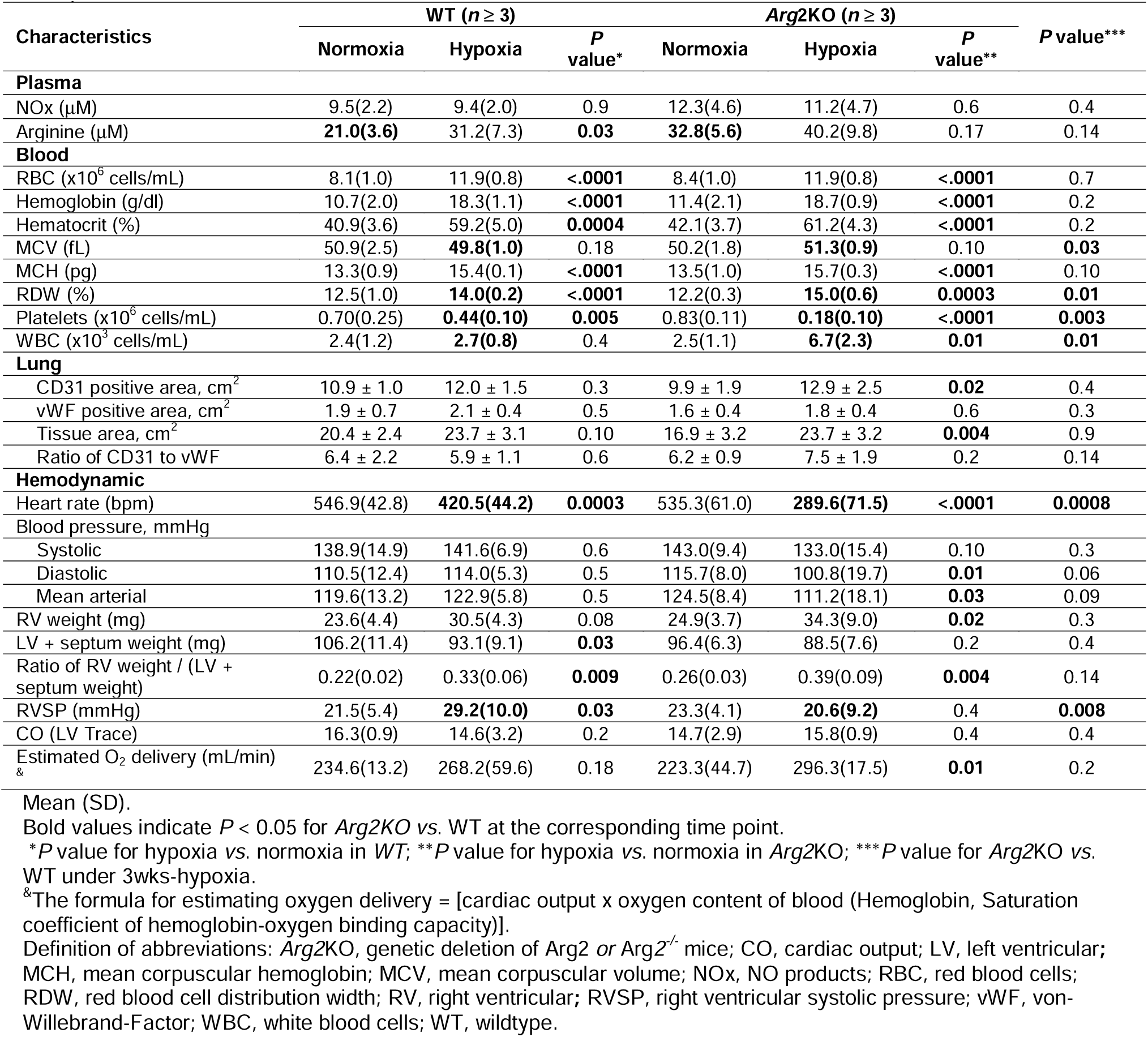
Hematologic and cardiovascular characteristics of *Arg*2KO and WT mice under 3 weeks long-term hypoxia or kept under normoxia.

| Characteristics | WT (*n* ≥ 3) |  |  | Arg2KO (*n* ≥ 3) |  |  | *P* value*** |
| --- | --- | --- | --- | --- | --- | --- | --- |
|  | Normoxia | Hypoxia | *P* value* | Normoxia | Hypoxia | *P* value** |  |
| **Plasma** |  |  |  |  |  |  |  |
| NOx (μM) | 9.5(2.2) | 9.4(2.0) | 0.9 | 12.3(4.6) | 11.2(4.7) | 0.6 | 0.4 |
| Arginine (μM) | **21.0(3.6)** | 31.2(7.3) | **0.03** | **32.8(5.6)** | 40.2(9.8) | 0.17 | 0.14 |
| **Blood** |  |  |  |  |  |  |  |
| RBC (x10<sup>6</sup> cells/mL) | 8.1(1.0) | 11.9(0.8) | **<.0001** | 8.4(1.0) | 11.9(0.8) | **<.0001** | 0.7 |
| Hemoglobin (g/dl) | 10.7(2.0) | 18.3(1.1) | **<.0001** | 11.4(2.1) | 18.7(0.9) | **<.0001** | 0.2 |
| Hematocrit (%) | 40.9(3.6) | 59.2(5.0) | **0.0004** | 42.1(3.7) | 61.2(4.3) | **<.0001** | 0.2 |
| MCV (fL) | 50.9(2.5) | **49.8(1.0)** | 0.18 | 50.2(1.8) | **51.3(0.9)** | 0.10 | **0.03** |
| MCH (pg) | 13.3(0.9) | 15.4(0.1) | **<.0001** | 13.5(1.0) | 15.7(0.3) | **<.0001** | 0.10 |
| RDW (%) | 12.5(1.0) | **14.0(0.2)** | **<.0001** | 12.2(0.3) | **15.0(0.6)** | **0.0003** | **0.01** |
| Platelets (x10<sup>6</sup> cells/mL) | 0.70(0.25) | **0.44(0.10)** | **0.005** | 0.83(0.11) | **0.18(0.10)** | **<.0001** | **0.003** |
| WBC (x10<sup>3</sup> cells/mL) | 2.4(1.2) | **2.7(0.8)** | 0.4 | 2.5(1.1) | **6.7(2.3)** | **0.01** | **0.01** |
| **Lung** |  |  |  |  |  |  |  |
| CD31 positive area, cm<sup>2</sup> | 10.9 ± 1.0 | 12.0 ± 1.5 | 0.3 | 9.9 ± 1.9 | 12.9 ± 2.5 | **0.02** | 0.4 |
| vWF positive area, cm<sup>2</sup> | 1.9 ± 0.7 | 2.1 ± 0.4 | 0.5 | 1.6 ± 0.4 | 1.8 ± 0.4 | 0.6 | 0.3 |
| Tissue area, cm<sup>2</sup> | 20.4 ± 2.4 | 23.7 ± 3.1 | 0.10 | 16.9 ± 3.2 | 23.7 ± 3.2 | **0.004** | 0.9 |
| Ratio of CD31 to vWF | 6.4 ± 2.2 | 5.9 ± 1.1 | 0.6 | 6.2 ± 0.9 | 7.5 ± 1.9 | 0.2 | 0.14 |
| **Hemodynamic** |  |  |  |  |  |  |  |
| Heart rate (bpm) | 546.9(42.8) | **420.5(44.2)** | **0.0003** | 535.3(61.0) | **289.6(71.5)** | **<.0001** | **0.0008** |
| Blood pressure, mmHg |  |  |  |  |  |  |  |
| Systolic | 138.9(14.9) | 141.6(6.9) | 0.6 | 143.0(9.4) | 133.0(15.4) | 0.10 | 0.3 |
| Diastolic | 110.5(12.4) | 114.0(5.3) | 0.5 | 115.7(8.0) | 100.8(19.7) | **0.01** | 0.06 |
| Mean arterial | 119.6(13.2) | 122.9(5.8) | 0.5 | 124.5(8.4) | 111.2(18.1) | **0.03** | 0.09 |
| RV weight (mg) | 23.6(4.4) | 30.5(4.3) | 0.08 | 24.9(3.7) | 34.3(9.0) | **0.02** | 0.3 |
| LV + septum weight (mg) | 106.2(11.4) | 93.1(9.1) | **0.03** | 96.4(6.3) | 88.5(7.6) | 0.2 | 0.4 |
| Ratio of RV weight / (LV + septum weight) | 0.22(0.02) | 0.33(0.06) | **0.009** | 0.26(0.03) | 0.39(0.09) | **0.004** | 0.14 |
| RVSP (mmHg) | 21.5(5.4) | **29.2(10.0)** | **0.03** | 23.3(4.1) | **20.6(9.2)** | 0.4 | **0.008** |
| CO (LV Trace) | 16.3(0.9) | 14.6(3.2) | 0.2 | 14.7(2.9) | 15.8(0.9) | 0.4 | 0.4 |
| Estimated O<sub>2</sub> delivery (mL/min) & | 234.6(13.2) | 268.2(59.6) | 0.18 | 223.3(44.7) | 296.3(17.5) | **0.01** | 0.2 |

Mean (SD).
Bold values indicate *P* < 0.05 for *Arg2KO* vs. WT at the corresponding time point.
\**P* value for hypoxia vs. normoxia in WT; \*\**P* value for hypoxia vs. normoxia in *Arg2KO*; \*\*\**P* value for *Arg2KO* vs. WT under 3wks-hypoxia.
<sup>&</sup>The formula for estimating oxygen delivery = [cardiac output x oxygen content of blood (Hemoglobin, Saturation coefficient of hemoglobin-oxygen binding capacity)].
Definition of abbreviations: *Arg2KO*, genetic deletion of *Arg2* or *Arg2*<sup>-/-</sup> mice; CO, cardiac output; LV, left ventricular; MCH, mean corpuscular hemoglobin; MCV, mean corpuscular volume; NOx, NO products; RBC, red blood cells; RDW, red blood cell distribution width; RV, right ventricular; RVSP, right ventricular systolic pressure; vWF, von-Willebrand-Factor; WBC, white blood cells; WT, wildtype.

*Arg*2KO and WT mice have similar red blood cells (RBC), hemoglobin, hematocrit, mean corpuscular volume (MCV) (Fig. 1C **–** F), and red blood cell distribution width (RDW) (all *P* > 0.05) (Table 1). Normal erythroid development shows a 1:2:4:8 ratio from stage I to stage IV in WT and *Arg*2KO mice (Fig. 2A, 2B) (31, 32). However, *Arg*2KO mice have lower erythroid progenitors than WT mice [(proerythroblast (stage I), *P* = 0.12; basophilic erythroblasts (stage II), *P* = 0.01; polychromatic erythroblasts (stage III), *P* = 0.005; orthochromatic erythroblasts (stage IV), *P* = 0.01) (Table 1) (Fig. 2B, 2D, 2E, 2H, 2I).

**Fig. 2.**
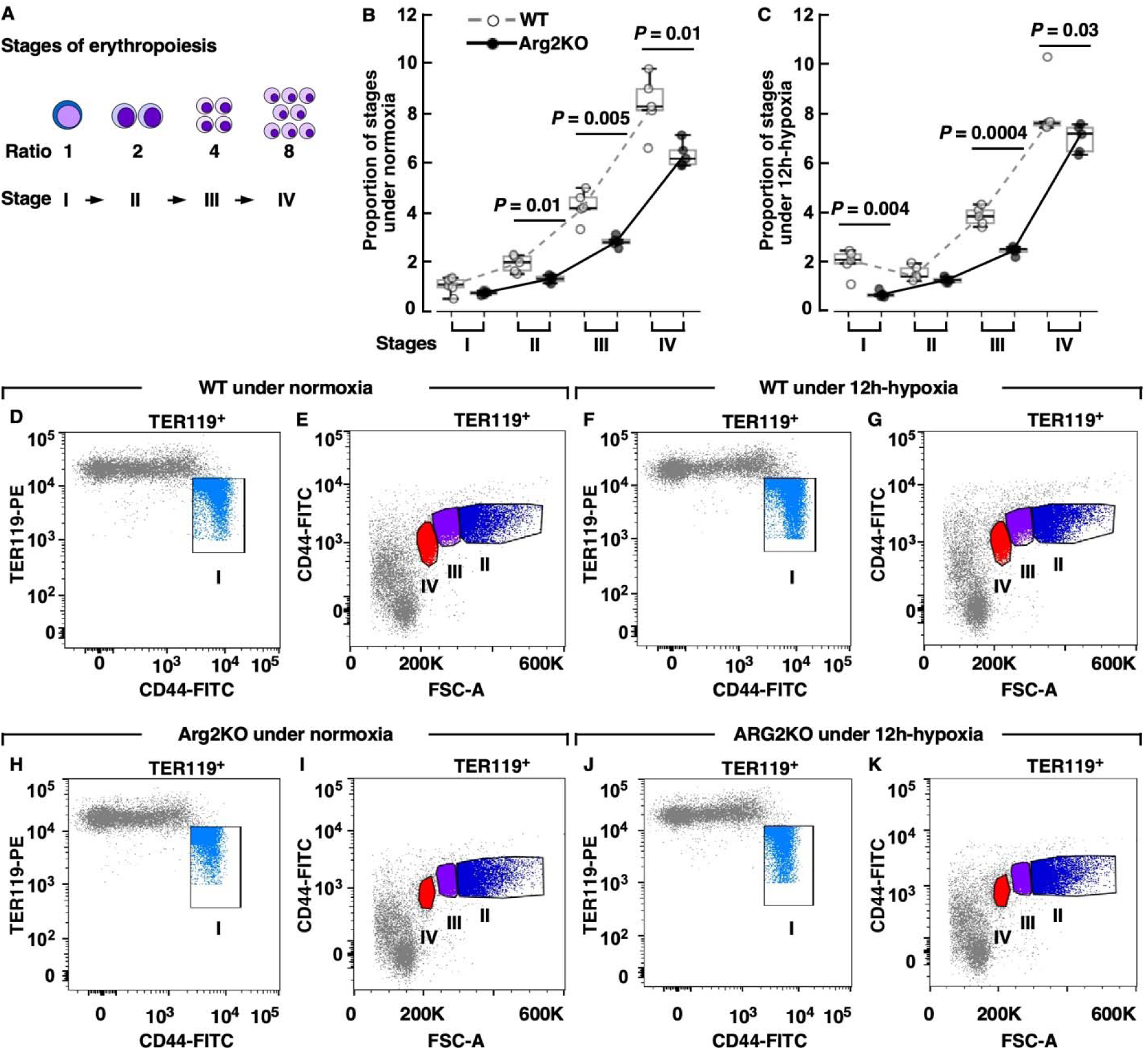
Normalized proportion of erythropoietic stage in *Arg2*KO and WT mice under normoxia or under 12hrs-hypoxia. [A] Normal erythroid development shows a 1:2:4:8 ratio from stage I to stage IV. **[B – C]** Normalized proportion of erythropoietic stage of stage I, stage II, stage III, and stage IV in *Arg2*KO and WT mice under normoxia (**B**) or under 12 hrs-hypoxia (**C**) (*n* ≥ 3 for each group). **[D – K]** Gating for erythroid progenitor stages measured via flow cytometry for WT mice under normoxia (**D and E**) or 12 h-hypoxia (**F and G**) and *Arg2*KO mice under normoxia (**H and I**) and 12 h-hypoxia (**J and K**). Proerythroblast (stage I), basophilic erythroblasts (stage lI), polychromatic erythroblasts (stage III), and orthochromatic erythroblasts (stage IV) were determined based on their expression of CD44, TER119, and size. All plots represent one million TER119+ cells. Ungated events are in grey. Proportion of erythropoietic stages are normalized to the proportion of stage I of WT mice under normoxia. TER-119 is a lineage marker for erythroid cells from early proerythroblast to mature erythrocyte stages in adult blood, spleen, and bone marrow. CD44 expressed on various cell types is used as a marker in flow cytometry to isolate specific cell populations in bone marrow.

Immunohistochemistry staining of endothelial markers CD31 and von Willebrand Factor (vWF) are used to assess vessels in lungs of *Arg2*KO and WT mice (33). As compared to WT mice, *Arg2*KO mice tend to have smaller lungs (total Lung Area cm^2^, *P* = 0.10) but with similar blood vessels (CD31 positive area cm^2^, *P* = 0.4; vWF positive area cm^2^, *P* = 0.4) (Table 2) (Supplementary Fig. 1A, 1C, 1E, 1G, 1I, 1K, 1M, 1O).

### Under short-term (6-, 12-, 48-, or 72-hours) hypoxia, *Arg*2KO have delayed and blunted erythropoiesis response

Plasma arginine and NO increase under hypoxia (Table 1) (Fig. 1A, 1B). NO levels peak at 12 hrs-hypoxia in WT, then drop quickly (ANOVA *P* = 0.003). In contrast, NO levels are significantly higher in *Arg*2KO than WT with long-lasting elevations up to 48 hrs-hypoxia (*P* = 0.03) (Table 1) (Fig. 1A). *Arg*2KO mice also have significantly higher arginine levels than WT mice under 6 hrs-, 12 hrs-, and 72 hrs-hypoxia (all *P* < 0.05) (Table 1) (Fig. 1B).

*Arg*2KO and WT mice significantly increase the variation in size among red blood cells under hypoxia (RDW, both ANOVA *P* < 0.05) (Table 1). RBC, hemoglobin, and hematocrit increase only in WT mice (all ANOVA *P* < 0.05) (Table 1) (Fig. 1C **–** E). Compared to WT mice, *Arg*2KO mice have significantly lower hematocrit and MCV under 6 hrs- and 72 hrs-hypoxia (Table 1) (Fig. 1E, 1F), suggesting *Arg*2KO have smaller red blood cells and are slower in response to short-term hypoxia.

Both *Arg*2KO and WT mice increase erythroid progenitor populations under hypoxia (all ANOVA *P* < 0.05) (Table 1). WT respond to 12 hrs-hypoxia with an increase in proerythroblast (stage I, *P* = 0.004), polychromatic erythroblasts (stage III, *P* = 0.0004) and orthochromatic erythroblasts (stage IV, *P* = 0.03) (Table 1) (Fig. 2C, 2F, 2G, 2J, 2K). At 48 hrs-hypoxia, *Arg*2KO increase the number of proerythroblasts (stage I, *P* = 0.0008) (Table 1). The proerythroblasts (stage I) in *Arg*2KO mice peak at 72 hours of hypoxia but tend to be lower than WT (*P* = 0.11) (Table 1). These data indicate that the erythropoiesis response to hypoxia in *Arg*2KO mice is delayed and blunted.

### Under 3 weeks long-term hypoxia, *Arg*2KO mice do not increase RVSP and have much lower heart rates

Under long-term hypoxia, *Arg*2KO and WT mice have similar plasma NO (*P* = 0.4). Plasma arginine levels in WT mice increase overall (*P* = 0.03) but tend to be lower than *Arg2*KO (*P* = 0.14) (Table 2).

Under long-term hypoxia, both *Arg*2KO and WT mice increase RBC, hemoglobin, hematocrit, MCH, and RDW (all *P* < 0.05) (Table 2). *Arg*2KO mice have much higher MCV and RDW as compared to WT mice (both *P* < 0.05) (Table 2), indicating that *Arg*2KO mice have larger mean RBC size and greater variation in RBC size.

Under long-term hypoxia, unlike WT, *Arg*2KO do not increase RVSP (*P* = 0.4) (Table 2) (Fig. 3A). WT and *Arg2*KO both increase ratio of RV weight / (LV + septum weight) (Table 2) (Fig. 3B **–** D), but *Arg2*KO have lower mean arterial blood pressures (*P* = 0.03), diastolic blood pressures (*P* = 0.01), and much lower heart rates than WT (*P* = 0.0008) (Table 2) (Fig. 3E **–** 3G). *Arg2*KO and WT mice have similar cardiac output (CO) (LV Trace) under normoxia and hypoxia (Table 2) (Fig.3H). The physiologic responses maintain oxygen delivery in WT mice. However, in *Arg2*KO, the changes lead to even greater oxygen delivery under hypoxia than in under normoxia (*P* = 0.01) (Table 2) (Fig. 3I),

**Fig. 3.**
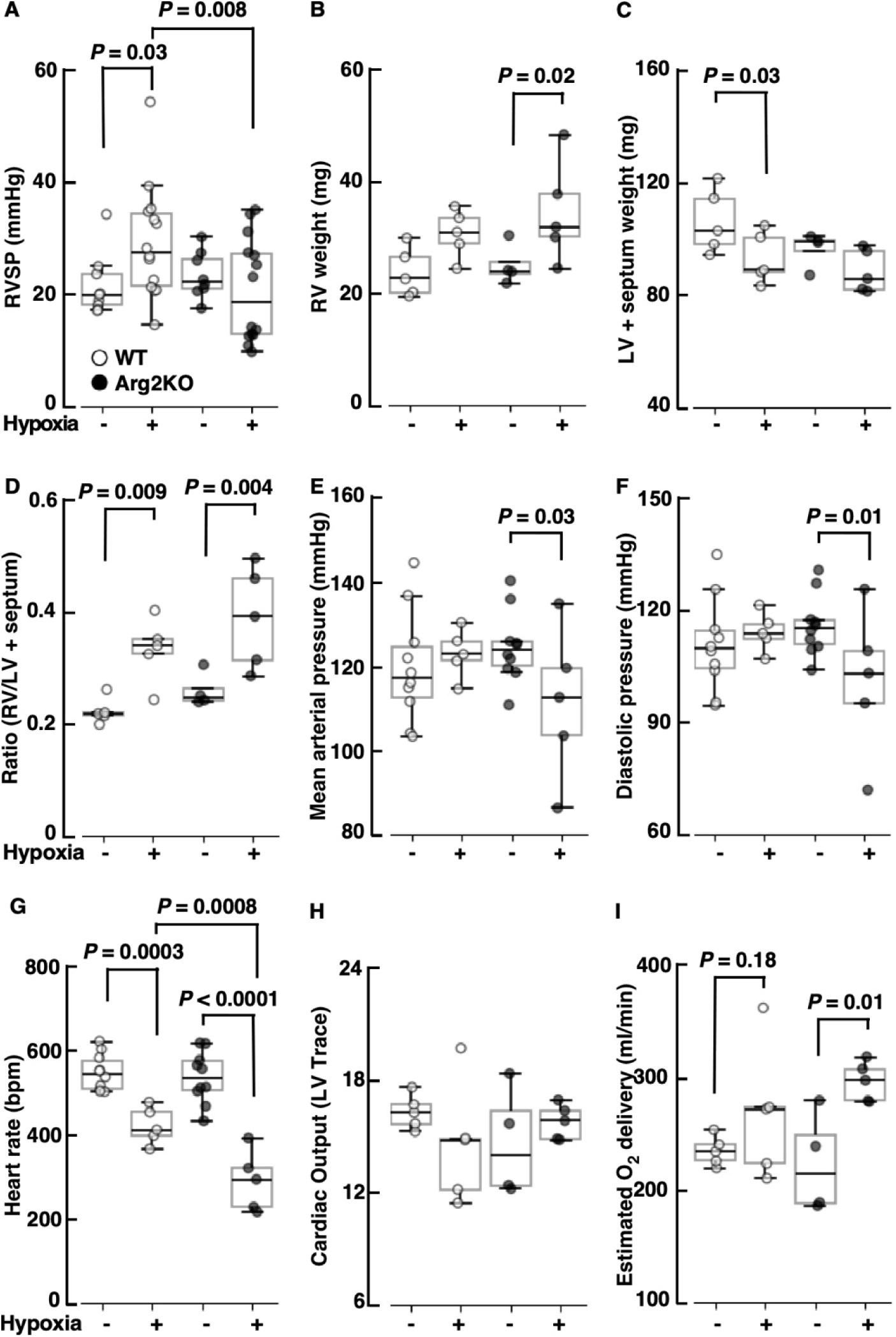
*Arg2*KO do not increase right ventricular systolic pressure (RVSP) under 3 weeks long-term hypoxia but have much lower heart rates. **[A]** Unlike WT (*n* ≥ 9), *Arg2*KO (*n* ≥ 8), did not increase RVSP under hypoxia. **[B – D]** WT (*n* = 5) had similar right ventricular (RV) weights under hypoxia (**B**), but lower left ventricular (LV) and septum weights (**C**) and increased ratio of RV / (LV + septum weight), while significantly increased weight of RV (**B**) and the ratio of RV / (LV + septum weight) (**D**) in *Arg2*KO (*n* ≥ 4) under hypoxia. [E – G] *Arg2*KO significantly decreased mean arterial blood pressure (**E**), diastolic blood pressure (**F**), and heart rate (**G**) under hypoxia (*n* ≥ 5). [H] *Arg2*KO (*n* ≥ 4) had similar cardiac output (LV Trace) compared with WT (*n* = 5) under hypoxia. **[I]** Compared with WT (*n* = 5), *Arg2*KO (*n* = 5) had significantly increased estimated oxygen delivery under hypoxia than under normoxia (*n* = 4).

Analysis of immunohistochemistry stains of CD31 and vWF shows that *Arg2*KO have larger lung size and CD31 positivity but not vWF positivity in lung tissue under long-term hypoxia compared to WT. Although both immunostains are used to identify endothelial cells, vWF, a glycoprotein, shows stronger staining in larger arteries and veins, while CD31, a transmembrane glycoprotein, is a more reliable marker to identify endothelial cells in lung tissue (33). *Arg2*KO mice increase lung size and CD31 positive blood vessels significantly under long-term hypoxia (total Lung Area cm^2^, WT *P* = 0.10, *Arg2*KO *P* = 0.004; CD31 positive area cm^2^, WT *P* = 0.3; *Arg2*KO *P* = 0.02) (Table 2), but not vWF positive blood vessels (vWF positive area cm^2^, WT *P* = 0.5, *Arg2*KO *P* = 0.3) (Table 2) (Supplementary Fig.1). The ratio of CD31 to vWF tends to be higher in lungs of *Arg2*KO mice than WT mice (*P* = 0.14) (Table 2). The findings suggest *Arg2*KO mice increase small vessel angiogenesis under long-term hypoxia (33). This might affect lung vasculature and pulmonary artery pressures and result in the greater oxygen delivery in *Arg2*KO (34, 35). Higher levels of NO in the *Arg2*KO mice might favor endothelial cell proliferation (16).

## Discussion

Genetic deletion of mitochondrial *Arg2* in mice significantly enhances oxygen delivery, primarily through cardiovascular adaptations. Under normoxic conditions, *Arg2*KO and WT mice exhibit similar RBC and hemoglobin concentrations, heart rates, and blood pressures. However, under short-term hypoxia, *Arg2*KO mice have a delayed and attenuated erythropoietic response compared to WT mice, and a sustained elevation of plasma arginine and NO levels. In chronic hypoxia, both *Arg2*KO and WT mice achieve similar RBC counts, but *Arg2*KO mice demonstrate notable cardiovascular protection. They do not experience an increase in RVSP, maintain lower mean systemic and diastolic blood pressures, and have significantly lower heart rates. Additionally, *Arg2*KO mice show a marked increase in small vessels in the lungs*, i.e.*, angiogenesis.

Hematopoietic (erythropoiesis) adaptations in some high-altitude populations, such as Andeans, are important to increase oxygen carrying capacity to survive lifelong hypoxic stress (1, 3, 6, 10, 11, 13, 15). Vascular adaptations are more prominent in high-altitude populations in Nepal and Ethiopia. Here, as previously reported (36), WT mice increase blood RBC, hemoglobin, hematocrit, and erythroid progenitor populations with acute hypoxia. The weak erythropoietic response to acute hypoxia in *Arg2*KO mice is a new observation. It is interesting to speculate that the dampened response is due to a change in intracellular arginine metabolism to spermidine (22). Spermidine is a key precursor in the biosynthesis of hypusine, a post-translational modification on the eukaryotic translation initiation factor 5A (eIF5A). In this process, a specific lysine residue on eIF5A is modified by addition of an aminobutyl from spermidine.

Hypusination of eIF5A is necessary for proper erythroid differentiation and red blood cell production. Impaired hypusination may occur in *Arg2*KO, which would impair erythropoiesis (22). Further studies are needed to determine the mechanisms.

Chronic hypoxia exposure is used as a model for pulmonary hypertension (28, 29, 37, 38). Mice with genetic deletion of eNOS have mild pulmonary hypertension under normoxia and an exaggerated pulmonary vasoconstrictive response to hypoxia (37, 38). Our group discovered that high-altitude natives who avoid hypoxic pulmonary hypertension have low levels of arginase activity and high NO, which are associated with greater blood flow and oxygen delivery to tissues. The greater vascular response is associated with lesser hematopoietic response in these populations (2, 3, 6, 7, 10, 15). In contrast to high-altitude populations, serum arginase activity is higher in patients with pulmonary arterial hypertension (PAH) as compared to healthy controls (21, 26).

Previous reports show that arginase inhibitors reduce right ventricular systolic pressure (RVSP), lung tissue remodeling, and improve NO bioavailability in models of pulmonary hypertension (39, 40). Oral arginine therapy lowers pulmonary artery systolic pressures in PAH patients with sickle cell disease (41). In this study, *Arg2*KO mitigates hypoxia-induced pulmonary hypertension. WT and *Arg2*KO both increase ratio of RV weight / (LV + septum weight), but *Arg2*KO do not have increases in RVSP. Furthermore, *Arg2*KO mice have a greater increase in lung size and vasculature, which suggests more angiogenic activity (34, 35).

Overall, this study shows that mitochondrial arginine metabolism modulates adaptive responses to hypoxia. Genetic deletion of *Arg2* results in higher NO levels, avoidance of pulmonary and systemic hypertension, and greater oxygen delivery under hypoxia, providing strong evidence that arginine metabolism regulates cardiovascular health.

## Author contributions

WX, KA, and SCE designed the research studies. KA, AJJ, EM, NW, DT, MN, and AM performed the experiments and acquired data. WX, KA, AJJ, SF and SCE analyzed the data. WX, KA, and SCE wrote the manuscript.

## Acknowledgments

The authors thank J. Peterson, A. Branicky, A. Helmick, and R. Zhang for help with the study. This work was supported by NIH grant HL060917. The authors thank the technical staff of the LRI Flow Cytometry Core for excellent assistance with instrument QC and setup. This work utilized a Sony ID7000 that was purchased with funding from the National Institutes of Health SIFAR grant S10OD025207.

Arg: arginine
Arg2KO: genetic deletion of Arg2 or Arg2-/- mice
CO: cardiac output
LV: left ventricular
MCH: mean corpuscular hemoglobin
MCV: mean corpuscular volume
NO: nitric oxide
NOx: NO products
RBC: red blood cells
RDW: red blood cell distribution width
RV: right ventricular
RVP: right ventricular pressure
RVSP: right ventricular systolic pressure
vWF: von Willebrand Factor
WBC: white blood cells
WT: wildtype

