## Supplementary Fig. 1 for "Hypoxia responses in arginase 2 deficient mice enhance cardiovascular health"

### Supplementary Figure and Legend

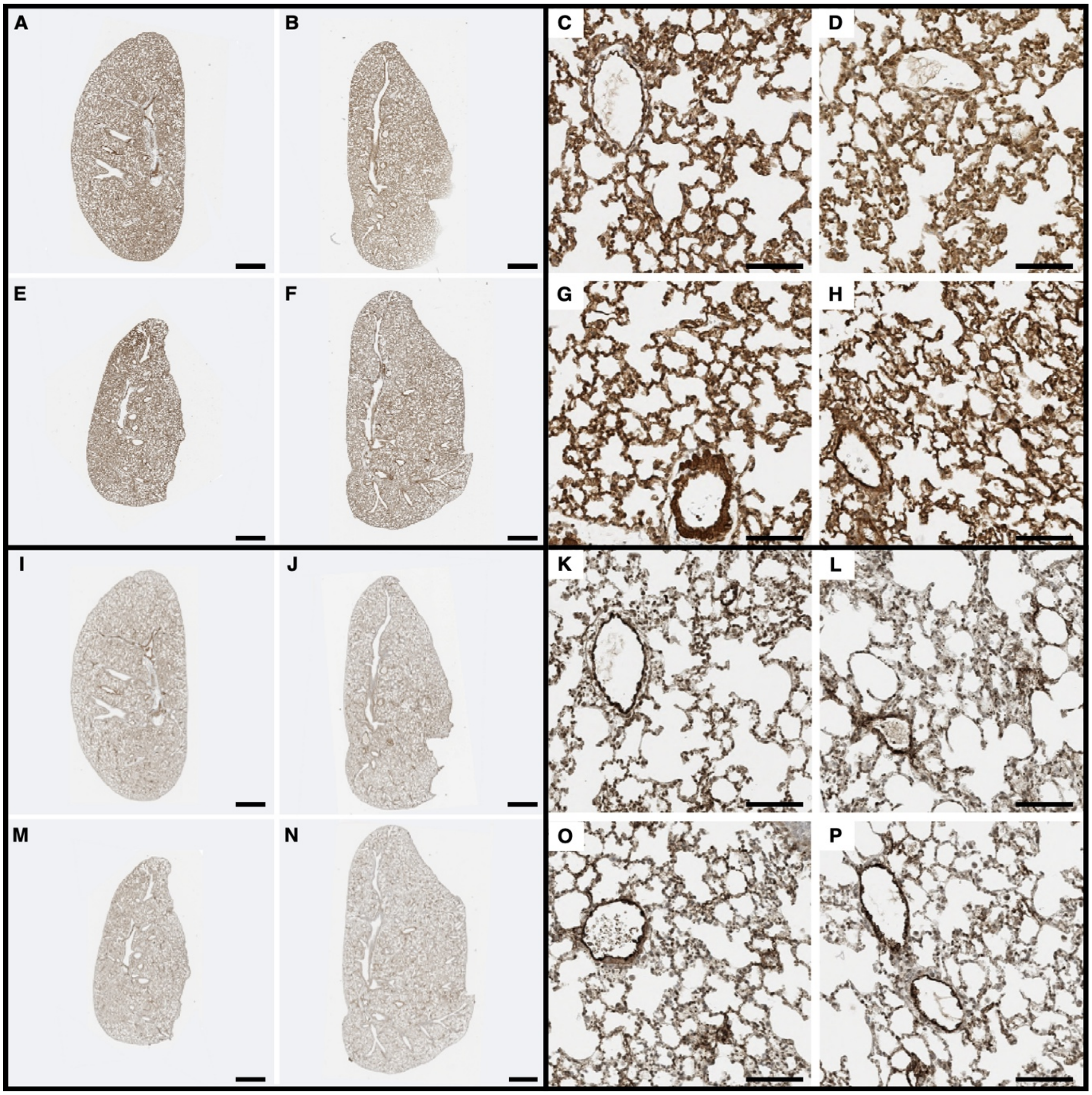


**Supplementary Fig. 1.** Immunohistochemical analyses of endothelial markers CD31 and von Willebrand Factor (vWF) in *Arg2*KO and WT mice lungs. **[A – H]** CD31 immunostains (brown) in *Arg2*KO **(E – H)** and WT mice lungs **(A – D)** under 3 weeks long-term hypoxia **(B, D, F, H)** or kept under normoxia **(A, C, E, G)**. CD31 staining in *C*, *D*, *G,* and *H* (scale bar: 1.2 mm) are high-power views of *A*, *B*, *E,* and *F* (Scale bar: 80 μm), respectively. **[I – P]** vWF immunostains (brown) in *Arg2*KO **(M – P)** and WT mice lungs **(I – L)** under 3 weeks long-term hypoxia **(J, L, N, P)** or kept under normoxia **(I, K, M, O)**. vWF staining in *K*, *L*, *O,* and *P* (scale bar: 1.2 mm) are high-power views of *I*, *J*, *M,* and *N* (scale bar: 80 μm), respectively. Images representative of sections from four *Arg2*KO and five WT lungs under normoxia, and five *Arg2*KO and five WT lungs under hypoxia.
